# A chromosome scale genome assembly and evaluation of mtDNA variation in the willow leaf beetle *Chrysomela aeneicollis*

**DOI:** 10.1101/2023.04.19.537531

**Authors:** Ryan R. Bracewell, Jonathon H. Stillman, Elizabeth P. Dahlhoff, Elliott Smeds, Kamalakar Chatla, Doris Bachtrog, Caroline Williams, Nathan E. Rank

## Abstract

The leaf beetle *Chrysomela aeneicollis* has a broad geographic range across Western North America, but is restricted to cool habitats at high elevations along the west coast. Central California populations occur only at high altitudes (2900-3450 m) where they are limited by reduced oxygen supply and recent drought conditions that are associated with climate change. Here we report a chromosome-scale genome assembly alongside a complete mitochondrial genome, and characterize differences among mitochondrial genomes along a latitudinal gradient over which beetles show substantial population structure and adaptation to fluctuating temperatures. Our scaffolded genome assembly consists of 21 linkage groups; one of which we identified as the X chromosome based on female/male whole genome sequencing coverage and orthology with *Tribolium castaneum*. We identified repetitive sequences in the genome and found them to be broadly distributed across all linkage groups. Using a reference transcriptome, we annotated a total of 12,586 protein coding genes. We also describe differences in putative secondary structures of mitochondrial RNA molecules, which may generate functional differences important in adaptation to harsh abiotic conditions. We document substitutions at mitochondrial tRNA molecules and substitutions and insertions in the 16S rRNA region that could affect intermolecular interactions with products from the nuclear genome. This first chromosome-level reference genome will enable genomic research in this important model organism for understanding the biological impacts of climate change on montane insects.

## INTRODUCTION

Breakthroughs in sequencing technology are now allowing us to produce high quality chromosome-scale genome assemblies for organisms across the tree of life. Beetles (order Coleoptera) constitute an exceptionally large group with more described species than any other insect order, yet their genomic resources lag behind other arthropods (Hotaling et al. 2021). This is surprising given that Coleoptera contains numerous well-studied species of significant economic and ecological importance. By broadening genomic resources in Coleoptera, we will begin to understand what has made this group one of the most successful animal radiations on earth and will deepen our knowledge of how they integrate into natural and agricultural systems.

The leaf beetle *Chrysomela aeneicollis* (Fig. 1) is found in montane and coastal habitats across Western North America (Brown 1956; Dellicour et al. 2014). It occurs at high elevation (2800-3300 m) in the Sierra Nevada range in California, where it feeds on willows in bogs and along streams(Smiley and Rank 1986). Decades of ecological and evolutionary studies have focused on populations found at high elevation (2800-3600 meters above sea level) in the southern Sierra Nevada in California (Dahlhoff and Rank 2000; Rank 1992; Rank et al. 2007; Smiley et al. 1985). These studies have revealed that Sierra Nevada populations of this insect are strongly dependent on annual fluctuations in temperature and snowpack (Dahlhoff et al. 2019; Roberts 2022), and show evidence of local adaptation to thermal microclimate and environmental oxygen (McMillan et al. 2005; Millstein 2006; Rank 1992; Rank et al. 2020).

**Figure 1.**
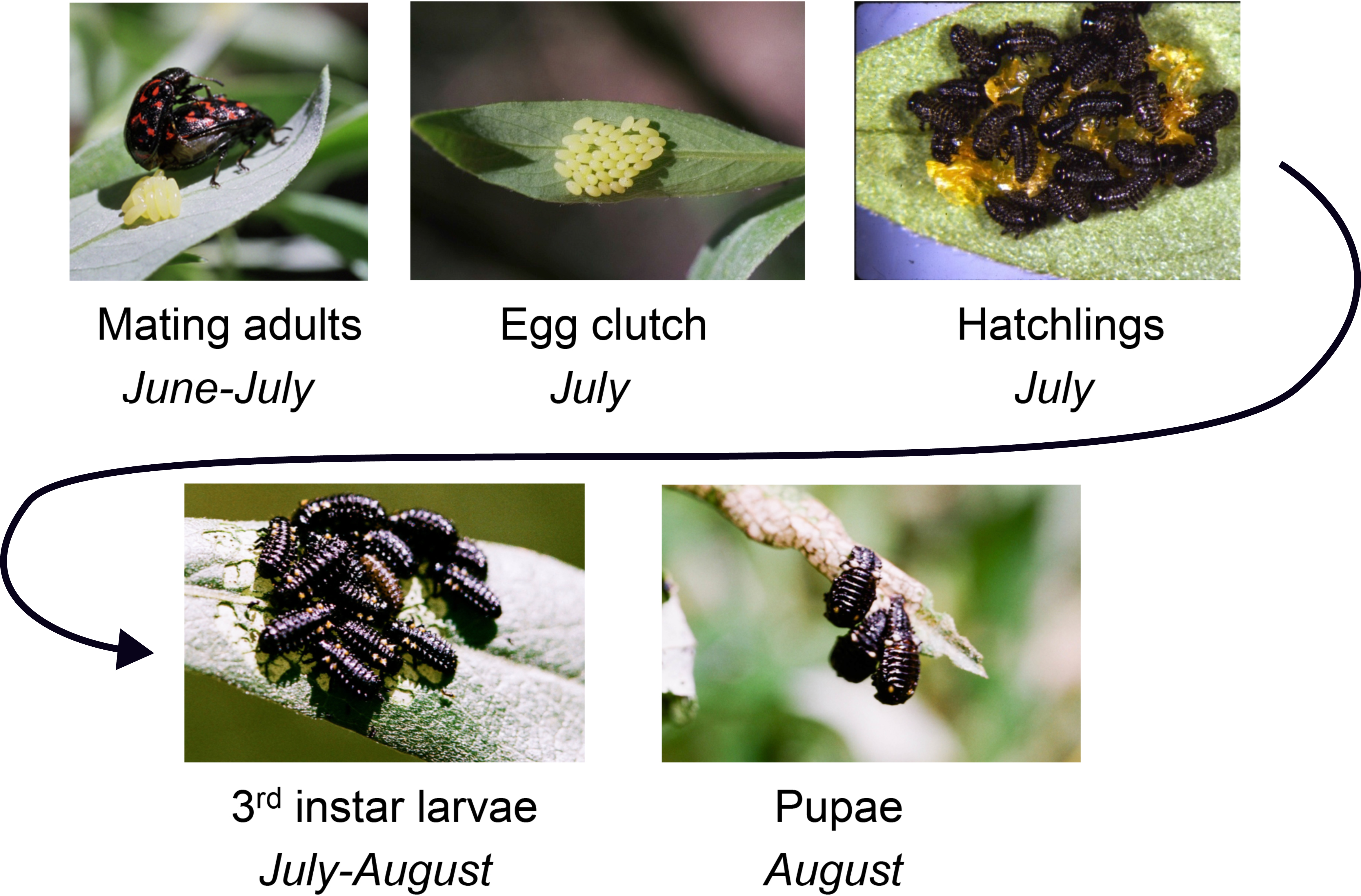
The willow leaf beetle *Chrysomela aeneicollis* (Schaeffer), a natural model system for evolutionary, physiological, and ecological genomics. Life cycle stages of this univoltine insect, which completes larval development during mid-summer.

Allele frequencies of key metabolic enzyme loci fluctuate, in a putatively adaptive way, with changes in climate (Dahlhoff et al. 2008; Rank and Dahlhoff 2002). Furthermore, recent evidence suggests that relationships between mitochondrial genotype, performance and reproductive success in Sierra Nevada populations depend on genotype at the nuclear enzyme locus *phosphoglucose isomerase* (Dahlhoff et al. 2019; Rank et al. 2020), providing a case study for the importance of mitonuclear epistasis in natural populations under field conditions (Camus 2020). The broader context for historical phylogeographic diversification of *C. aeneicollis* populations in western North America has been described for both mitochondrial and nuclear loci (Dellicour et al. 2014). Together with extensive ecological data on host plant relationships and food web dynamics (Otto et al. 2008; Rank 1994; Smiley et al. 1985), these studies provide a unique understanding of the natural history, physiology and evolutionary ecology of an insect species in its natural context.

Here, we report both the nuclear and mitochondrial genome of *Chrysomela aeneicollis*. This is the first genome assembly for this species. We also provide careful annotations of transcribed protein-coding and RNA gene products of mitochondria. In addition, we assess differences in mitochondria of 12 individuals sampled from three high altitude populations in the Eastern Sierra Nevada (Rock Creek, Bishop Creek, and Big Pine Creek). Finally, we identify potential secondary structure differences in mitochondrial rRNA and tRNA molecules in these populations and discuss the role these variants may play in mechanisms of local adaptation to environmental temperature and oxygen levels.

## MATERIALS AND METHODS

### Initial assembly with 10x Genomics

A single adult female beetle from Mosquito Flat in Rock Creek (37° 26′ 29.6” N, 118° 44′ 48.3” W; 3067 meters ASL) was collected in September 2017, kept alive in the laboratory until Feb 2018, then frozen at −80°C until DNA extraction. High-molecular weight genomic DNA was extracted by staff of the University of California Davis DNA Technologies and Expression Analysis Core Laboratory (Davis, CA USA). Blue Pippin size selection was used to retain fragments > 40 kb and resulting DNA was used to construct sequencing linked-read libraries using 10x Genomics Technology. PippinHT size selection of the final libraries (350-650bp) was used to maximize read 2 quality when sequenced using an Illumina HiSeqX10 at Novogene (Sacramento, CA USA). The resulting sequence data (150 bp PE) was assembled into a draft genome using Supernova v2.0.0 (Weisenfeld et al. 2017).

Assemblies were made using 56x coverage (224M reads, following “optimal” amount of data recommendations) and 99x coverage (395M reads, all the sequence data). For each assembly, Supernova output was generated in three styles (Pseudohap, Megabubble and Raw), though only the Pseudohap output, which generates a single record per scaffold, was used for subsequent analysis. We used BUSCO v.2.0 (Simão et al. 2015) on CyVerse with three references (Metazoa, Arthropoda and Insecta_odb9) to determine which assembly was more complete.

### Hi-C scaffolding

To further scaffold the Supernova-based assembly, we used chromatin conformation capture to generate Hi-C data (Lieberman-Aiden et al. 2009). Hi-C libraries were generated as outlined in (Bracewell et al. 2019) using a Dnase digestion method (Ramani et al. 2016) and a single female adult as input. The resulting DNA library was prepared using an Illumina TruSeq Nano library prep kit and was sequenced on a HiSeq 4000 with 100bp PE reads. We used the Juicer pipeline (Durand et al. 2016b) to map Hi-C reads and generate contact maps based on 3D interactions. We then used the 3D-DNA pipeline (Dudchenko et al. 2017) to orient and place contigs. 3D-DNA output files were visualized and checked for accuracy using Juicebox (Durand et al. 2016a) with verification and modifications to scaffolding done using built-in tools. The final assembly was scaffolded together with 500 Ns between each contig.

### Identifying the putative X chromosome scaffold

We used whole genome sequencing data of sexed individuals (described below in mtDNA section) to identify the putative X chromosome scaffold. In male heterogametic taxa (most Coleoptera), females are expected to have roughly double the sequencing coverage of a male over the X chromosome. Therefore, we mapped Illumina 150bp PE reads to our draft genome using BWA MEM (Li and Durbin 2009) and compared normalized female/male coverage over all linkage groups using bedtools2 (Quinlan and Hall 2010).

### Repetitive element identification and genome masking

We used EDTA (Ou et al. 2019) and RepeatModeler2 (Flynn et al. 2020), to identify repetitive sequences in the genome assembly. With these repeat libraries, we then masked the genome with RepeatMasker version 4.0.7 using the -no_is and -nolow flags. To characterize the proportion of sequence that was identified as repetitive and the genomic distribution of specific repetitive elements, we used bedtools2 (Quinlan and Hall 2010). We identified repeat-enriched regions of the genome as those areas where at least two consecutive genomic intervals (i.e., 100 kb region given 50 kb windows) were estimated to be at least double the genome-wide average.

### Transcriptome data

To generate a transcriptome to aid in genome annotation (below), we used available RNA-seq data from whole bodies of third instar larvae from an experiment investigating insect performance and gene expression (Elmore 2021). We chose data from three larvae, one each from a different experimental treatment (A73, A220, A223), in an attempt to build a diverse transcript library.

### Genome annotation

To annotate the assembly, we used the repeat-masked genome and the MAKER2 annotation pipeline (Holt and Yandell 2011) to identify gene models. The *ab initio* gene predictors SNAP (Korf 2004) and AUGUSTUS (Stanke and Waack 2003) were used to guide the annotation with the protein set from *Tribolium castaneum*. We also cleaned RNA-seq raw reads using Seqyclean (https://github.com/ibest/seqyclean) and then used HISAT2 (Kim et al. 2019) and the -dta flag to align the reads to the draft genome. We used StringTie2 (Kovaka et al. 2019) to build a *C. aeneicollis* transcriptome for use as a guide in annotation. Input transcripts used for annotation were created from the transcriptome file using gffread. To compare our final MAKER2 annotation to the model beetle *Tribolium castaneum* annotation, we used orthoDB (Kriventseva et al. 2018) and translated protein sequences from the two species (version Tcas_5.2) and compared them locally.

### Final genome assembly cleaning

As a final cleaning step, we re-examined small sequences that were not incorporated into Hi-C scaffolds. These sequences were screened to determine if they were non-target sequences (i.e., mtDNA, *Wolbachia*, or *Spiroplasma*) or contained annotated protein coding genes. We also removed unscaffolded contigs that were not found to have any protein coding genes.

### mtDNA assembly and analysis

We used the mitochondrial genome program NOVOPLASTY to assemble mitochondrial genomes of 12 *C. aeneicollis* individuals, four each from Rock Creek, Bishop Creek, and Big Pine Creek. These individuals had been sampled as first instar larvae and reared to third instar under common garden conditions at 3000 m ASL during summer 2015 (Smeds 2022). Before extraction, larvae were dissected, flash-frozen in liquid nitrogen and stored at −80C until DNA extractions were performed in the laboratory of G. Lanzaro (University of California-Davis). Individual larval abdomens were placed in 2.0 mL microcentrifuge tubes along with 100mL of MagMAX PK enzyme mix (2mL of 100mg/mL Proteinase K and 98mL of PK buffer, both from Thermofisher) and a 3mm diameter steel bead, then homogenized at 30Hz for 30s in a Qiagen Tissuelyser. Samples were then centrifuged at 10,000 *g* for 1 min in a benchtop microcentrifuge followed by 2 h incubation at 56°C in a shaking incubator. After incubation, samples were centrifuged and transferred to a 96-well plate with 100µL of MagMAX DNA lysis buffer, and incubated at room temperature for 10 min. A solution containing 215µL of MagAttract mix (100µL Buffer AL, 100mL isopropanol, and 15µL MagAttract Suspension) was added to each well, and the plate was loaded into a Qiagen BioSprint 96 extraction machine. DNA was extracted using BioSprint DNA Blood Kit reagents and BS96 DNA Tissue protocol. We also assembled the mitochondrial genome for a closely related species using one *Chrysomela lapponica* individual collected in Fjellas, Norway (69° 18’ 15’’ N; 19° 31’ 03’’ E) in June 2012 and stored at −80 °C until DNA extraction in 2019 using a Qiagen DNAEasy animal tissue kit.

We constructed genomic libraries for *C. aeneicollis* and *C. lapponica* using a Nextera DNA Flex kit with Nextera DNA CD indexes (Illumina). Each library was constructed with 500ng of starting material. We quantified input DNA preps and finished libraries on a Qubit 3.0 Fluorometer using dsDNA high sensitivity reagents and assessed library quality using an Agilent 2100 Bioanalyzer with dsDNA high sensitivity chip. 150bp PE sequencing was performed by Novogene (Davis, California) on an Illumina Hiseq X Ten, with twelve *C. aeneicollis* libraries sequenced on one lane and one *C. lapponica* library on a second lane.

We used Novoplasty 4.0 to assemble mitochondrial genomes. Before running Novoplasty, we used the perl script ‘filter_reads.pl’ to extract reads with putative mitochondrial sequence. When we ran Novoplasty, we used *cytochrome oxidase I* sequence from a prior draft genome, which was also compared to Sanger sequences obtained from prior studies (Rank et al. 2020) as seed sequence, and used default settings in the Novoplasty config.txt file for initial assembly. To annotate genomes of *C. aeneicollis* and *C. lapponica*, we used the program MITOS (http://mitos.bioinf.uni-leipzig.de/index.py). After initial automated annotation, we manually compared gene lengths and positions among both species and to a complete mitochondrial genome of the related species *Gonioctena intermedia* (Coleoptera: Chrysomelidae; accession number KX922881.1). Based on recommendations made by (Cameron 2014), we extended protein coding genes to their corresponding stop codons, allowing for more overlapping genes than the automated output provided. Once we completed these annotations, we validated nucleotide and amino acid sequences for 13 protein-coding genes and ribosomal RNA regions using BLAST and comparisons to relatives of *C. aeneicollis*. Annotations for tRNA coding genes were validated using the web-based tRNAscan-SE program (Chan and Lowe 2019). This application was also used to compare predicted secondary structure configurations for variants at the tRNA-Leu2 and tRNA-Lys regions. Predicted secondary structures for rRNA variants were analyzed and visualized using the web-based Vienna RNA Websuite (Lorenz et al. 2011). Once the genome was annotated, we removed nucleotides corresponding to the AT-rich region from genomes before final alignment and comparison of genetic variants. The alignments of the transcribed region and annotations for the 12 sequenced individuals were made using the genomics software platform Geneious (version 9.1.8), which we also used to generate the VCF file.

To understand the evolution of mtDNA in *C. aeneicollis*, we inferred relationships by using Maximum Likelihood and a General Time Reversible model with 1000 bootstrap replicates in MEGA-X (Kumar et al. 2018). Branch lengths were measured in the number of substitutions per site. This analysis involved 13 nucleotide sequences with a total of 14,622 positions in the final dataset. The alignment is in supplementary materials. All plots of relevant nuclear and genomic features were plotted using the *circlize* (Gu et al. 2014) and *karyploteR* (Gel and Serra 2017) packages in R.

## RESULTS

Our initial assembly using 56x (Assembly 1) and 99x (Assembly 2) coverage data produced 31,942 and 37,551 scaffolds, respectively. The total length of Assembly 1 was 475,329,771 bp and the length of Assembly 2 was 536,018,598 bp. The scaffold N50 for Assembly 1 was 1,120,830 and was 1,107,799 for Assembly 2. To determine which genome assembly to move forward with for the Hi-C scaffolding, we compared BUSCO results between the two and found them to be similar, with the number of complete BUSCOs higher, and with less missing, in Assembly 1 (Assembly 1: C:95.0%[S:91.6%,D:3.4%],F:3.6%,M:1.4%,n:1066. Assembly 2: C:94.7%[S:92.0%,D:2.7%],F:3.4%,M:1.9%,n:1066). Given these results, and assembly software recommendations, we moved forward with Assembly 1.

Our Hi-C data allowed us to scaffold Assembly 1 and we confidently identified 21 total linkage groups ranging in length from 24,271,041 bp to 7,025,367 bp (Fig. 2A and Table 1). Whole genome sequencing coverage differences between a female and male allowed us to identify the X chromosome (Fig. 2B). The karyotype is unknown for *C. aeneicollis* and therefore it is currently unclear if these linkage groups represent entire chromosomes or are individual arms of metacentric chromosomes. Some scaffolds likely represent complete metacentric chromosomes given clear associations on the diagonal of what would appear to be two chromosomal arms (Fig. 2A; LG4, LG5 and LG11). This Hi-C contact pattern is similar to contact maps of metacentric chromosomes in Drosophila (Bracewell et al. 2019; Bracewell et al. 2020). In addition to 21 linkage groups, a number of contigs and scaffolds were unable to be placed into linkage groups but were retained for additional downstream filtering.

**Figure 2.**
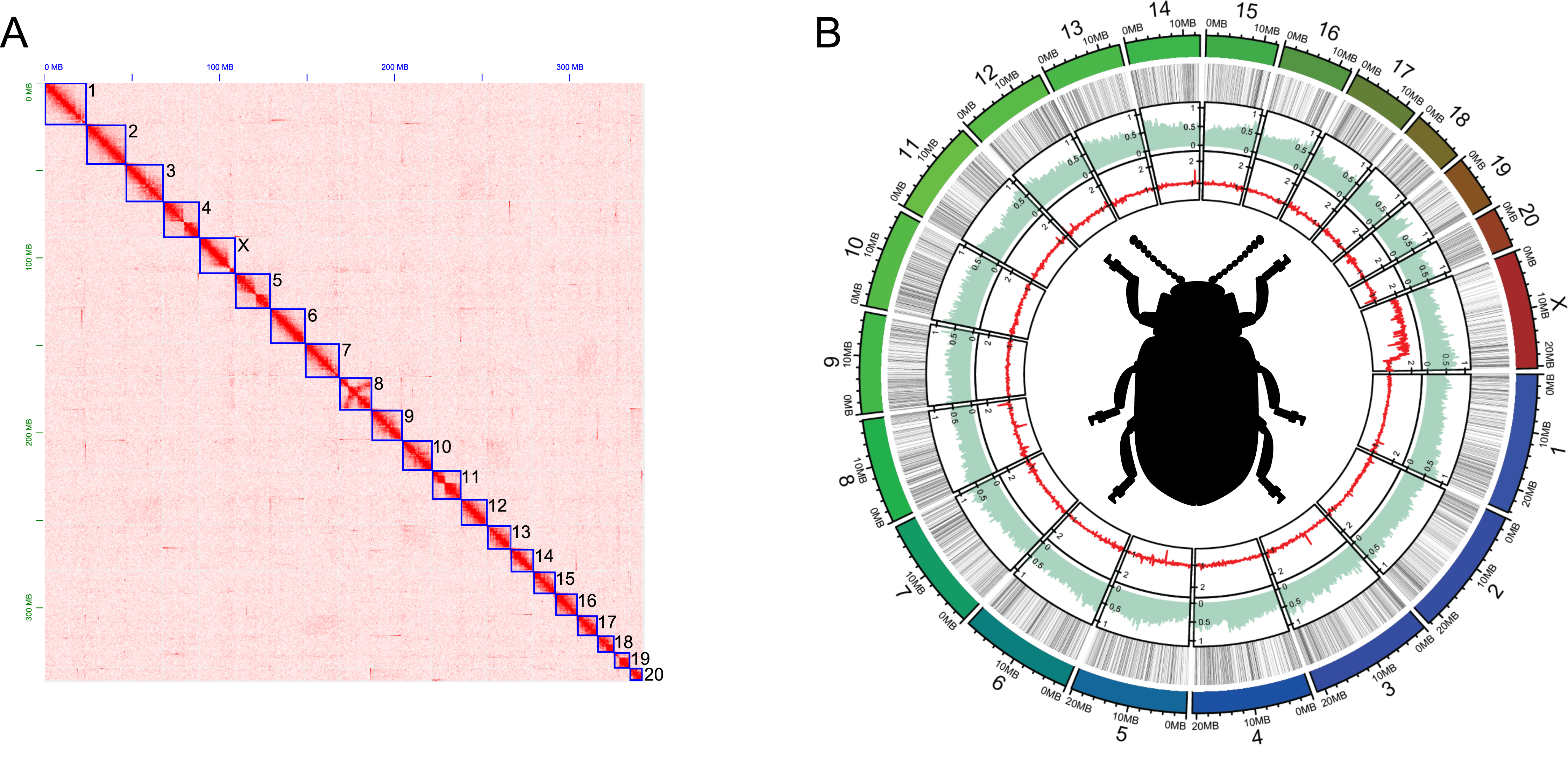
Nuclear genome structure. A) Hi-C contact heatmap showing chromosome level scaffolding of the 10X assembly. Twenty-one total linkage groups were found (shown in blue boxes). The observed bowtie patterns in the contact heatmap for some scaffolds (e.g., 4, 5, 11) are consistent with metacentric chromosomes. B) Circular plot showing the draft genome assembly and from the outermost track moving inwards a) the lengths of the 21 linkage groups b) a heatmap showing low (white) to high (black) density of scaffolding Ns, c) the percentage of sequence identified as repetitive (50 kb windows) and d) the normalized female/male whole genome sequencing coverage (red).

Repetitive sequences (transposable elements and tandemly repeated satellite sequence) were found to make up 43.5% of the genome assembly (Fig. 2B). We did not observe any enrichment in repetitive elements across LGs when all repeat families were grouped together (Fig. 2B). However, we found that specific repeat families were more frequently encountered than others, and LINEs constitute 7.1% of masked bases in the genome, LTR elements make up 5.4%, and DNA elements 3.8%. We found 26.2% of masked bases were unclassified and await future work. We found an increase in LTRs on the ends of most scaffolds suggesting a non-random distribution of these repeats that hints to their enrichment near centromeric or telomeric regions (Fig. S1).

The cleaning of raw RNAseq reads from three individuals left us with a total of 21M, 25M, 21M pairs of reads, and our combined transcriptome identified a total of 38,792 transcripts that were then used as evidence during annotation. Maker2 identified a total of 12,586 gene models in *C. aeneicollis,* with the number of protein coding genes on the 21 linkage groups ranging from 202 - 678 (Table 1). Using OrthoDB, we were able to identify a total of 8,187 orthologous genes with *Tribolium castaneum* (Herndon et al. 2020) thus providing some functional characterization of the *C. aeneicollis* annotation for future exploration. As a final cleaning step, we removed all contigs and scaffolds where the top two BLAST hits were either *Wolbachia* or *Spiroplasma* (known endosymbionts), the contig/scaffold did not have at least one annotated protein coding gene from above, or the contig/scaffold was a duplicated sequence already found in a linkage group or another scaffold/contig. After these filters, we retained 2,015 unplaced scaffolds (contigs) that contained 2,263 genes (Table 1). The above filters removed at total of 121.4 Mb of sequence. We visualized gene density across all LGs and any potential negative correlation with repeat density (Fig. S1). For some linkage groups, we observed a general trend where one end of an LG was gene poor and showed increased repetitiveness, which is indicative of a telocentric or subtelocentric chromosome (e.g., LG15 and LGX). However, most LGs showed no obvious spatial patterns of gene enrichment/depletion, suggesting genes are fairly evenly distributed across most LGs.

A query of genes involved in the Electron Transport Chain pathway of cellular respiration based on KEGG identifiers revealed that 59% of genes found in the published *Tribolium castaneum* genome were found in our *C. aeneicollis* assembly (41 of 69 genes, Table S2). The assembly also includes the full coding sequence for the nuclear metabolic enzyme *phosphoglucose isomerase* (PGI), which is located on LG3. Preliminary results indicate that the PGI electromorphs that have been investigated extensively in prior studies (Dahlhoff and Rank 2000; Rank et al. 2007) are determined by a non-synonymous substitution located in the first exon of the PGI coding sequence (Smeds 2022).

The reference sequence for the *C. aeneicollis* mitochondrion was generated in Novoplasty as a circularized genome of 18,007 bp. In the reference individual, the control region had a length of 3,390 bp, with 14,617 bp remaining for the transcribed region. For eight *C. aeneicollis* individuals and the *Chrysomela lapponica* individual, Novoplasty also assembled a circular genome. For four individuals, Novoplasty was only able to assemble linear contigs with multiple contigs in the control region. Sizes of *C. aeneicollis* mt genomes ranged from 17,158 to 18,794 bp (mean = 17,882.2, S.D. = 521.8, *n* = 8); the mt genome of the *Chrysomela lapponica* individual was somewhat smaller (16,750 bp). The transcribed region differed among individuals by only two base pairs and therefore the observed variation in mitochondrial genome size is largely due to differences in control region size rather than size of the transcribed region. One individual from Bishop Creek had an extra A nucleotide in the tRNA^Gln^ molecule, and six individuals (four from Big Pine Creek and two from Bishop Creek possessed a 2 bp insertion in the large ribosomal subunit. The final annotations for the reference sequence revealed seventeen cases of overlapping gene products within the transcribed region. Seven cases involved genes that overlapped with more than one other gene; eight between protein coding genes and tRNA gene products, four among protein coding genes alone and four tRNA gene regions. One was found between the small ribosomal subunit and a tRNA molecule.

The mitochondrial genomes of 12 sampled *C. aeneicollis* individuals (Fig. 3) showed substantial variation in the transcribed region, with nine distinct haplotypes recognized and four individuals (three from Big Pine Creek and one from Bishop Creek) sharing an identical DNA sequence (Fig. 4A, Table S1 for pairwise percent sequence divergence). Two distinct groups of haplotypes can be easily recognized; half of Bishop Creek individuals clustered together with individuals from the northern drainage Rock Creek and half clustered with the southern drainage Big Pine Creek (Fig. 4B). Fifty-three nucleotide differences occurred between RC64 and four individuals, all from Big Pine Creek, and the minimum number of differences between Rock Creek and Big Pine Creek individuals was forty-nine. This results in a substantial level of differentiation among northern and southern populations. The number of substitutions separating *C. lapponica* from *C. aeneicollis* was considerably greater (ranging from 253 to 260), indicating that the North American *C. aeneicollis* is monophyletic compared to the European *C. lapponica*.

**Figure 3.**
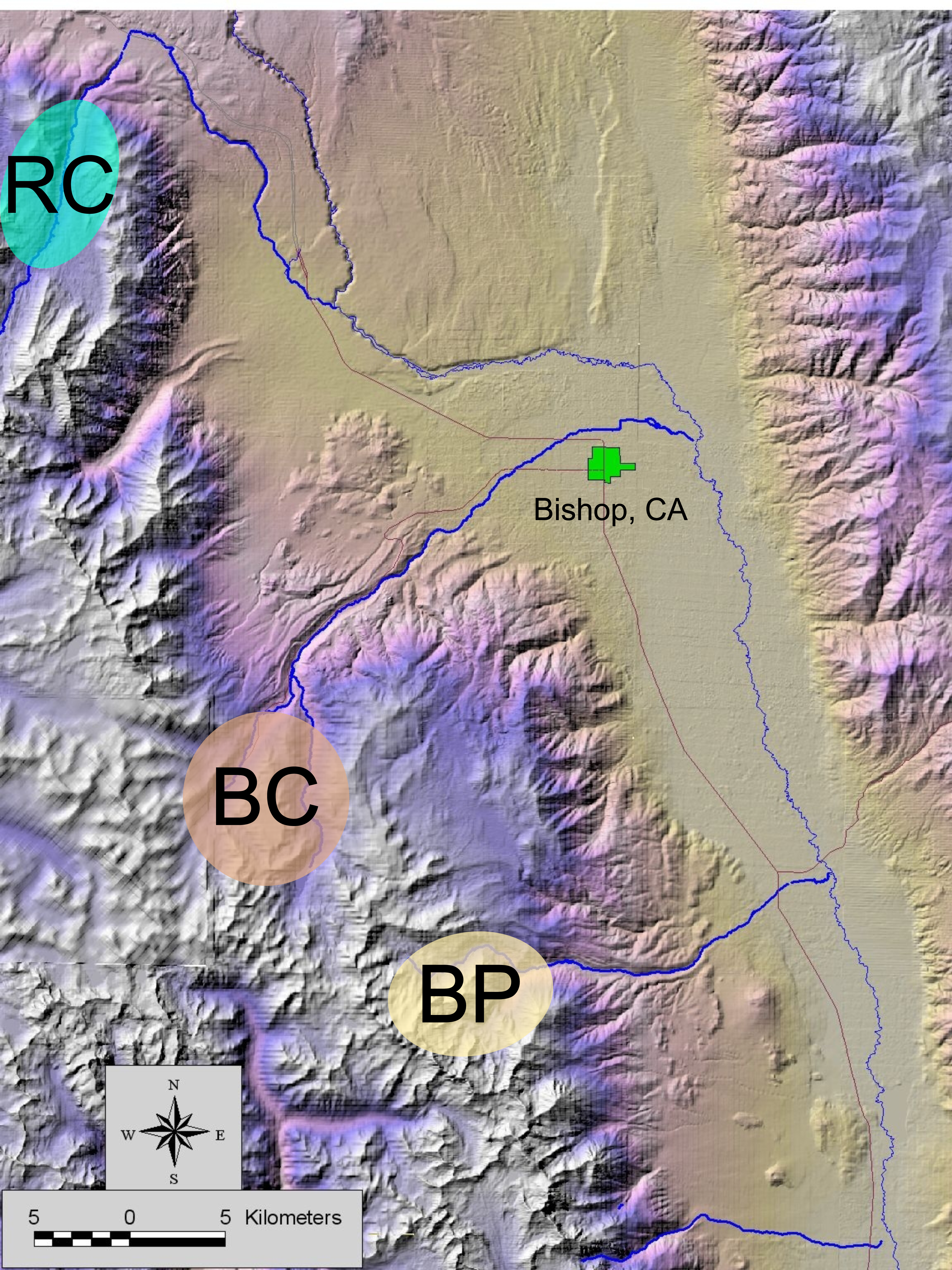
Collection locations of 12 *Chrysomela aenecollis* used in whole genome resequencing for mtDNA characterization. Individuals were initially collected from three distinct drainages identified as Rock Creek (RC), Bishop Creek (BC) and Big Pine (BP).

**Figure 4.**
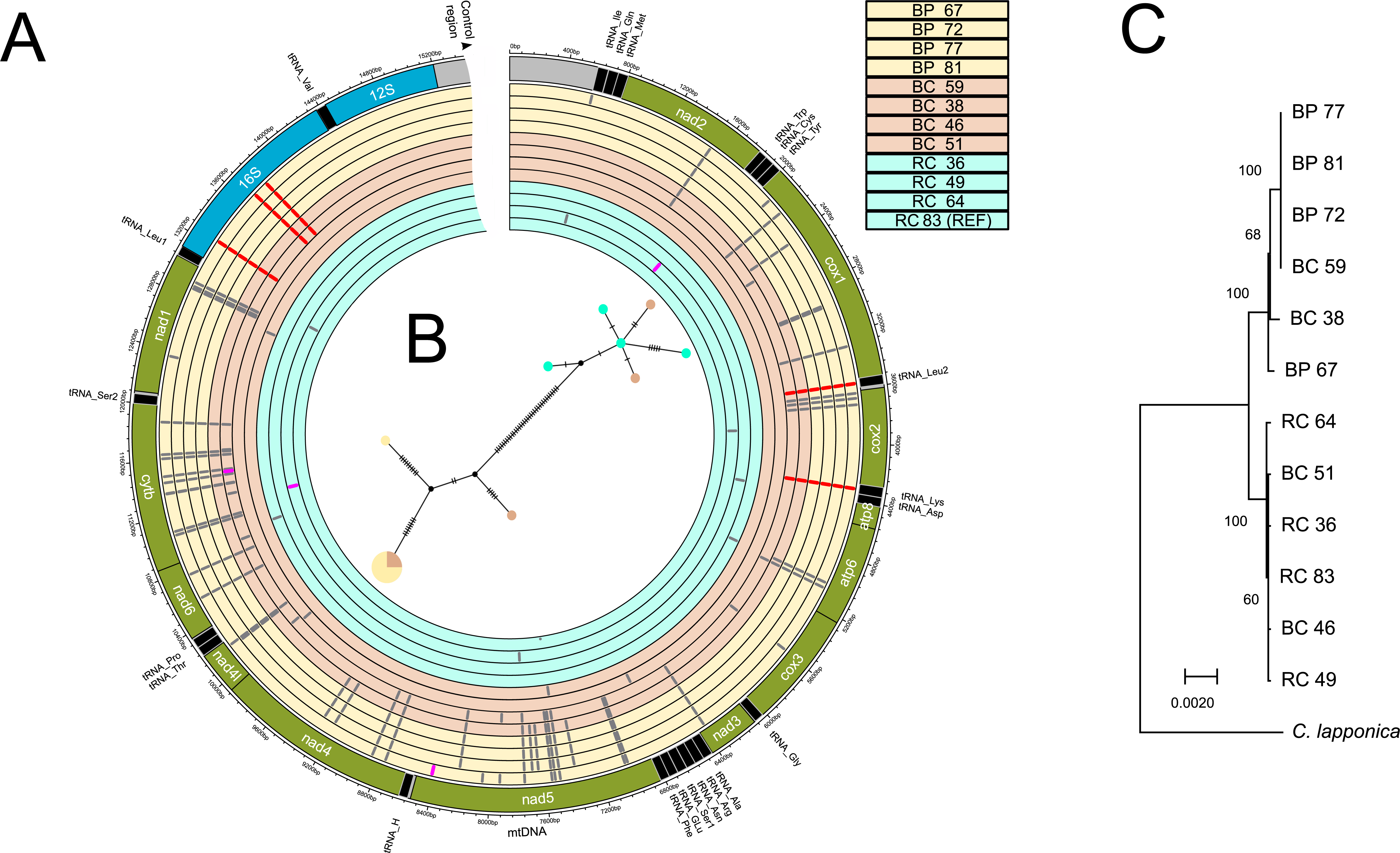
Mitochondrial genome structure. A) Circular representation of the assembled mtDNA genome from *C. aenecollis* (outer track) with the locations of tRNAs (black), protein coding genes (green), and rRNA (blue). The control region is not shown in its entirety. The inner twelve tracks show variants found in 4 individuals each from Big Pine (BP), Bishop Creek (BC) and Rock Creek (RC). An individual from Rock Creek was used as the reference (RC_83) and is the innermost track. Variants found in each mtDNA genome are shown as tick marks in their respective location. Grey marks show variants in tRNAs or protein coding genes that were synonymous or did not influence tRNA structure. Magenta marks are locations of nonsynonymous variants. Red tick marks highlight variants that were found in tRNAs or rRNA and appear to alter the secondary structure of the molecule. B) Haplotype network showing the relationships and number of nucleotide differences (black ticks) between mitotypes. C) Maximum likelihood phylogeny showing evolutionary relationships between mtDNA from 12 individuals with *C. lapponica* as an outgroup. Bootstrap values ≥ 60 are shown.

| Linkage group | length (bp) | Protein Coding |  | N (%) | Repetitive (%) |
| --- | --- | --- | --- | --- | --- |
|  |  | Genes | GC (%) |  |  |
| Scaf_01 | 24,271,041 | 678 | 35.0% | 15.9% | 42.8% |
| Scaf_02 | 22,353,350 | 632 | 34.9% | 15.2% | 41.7% |
| Scaf_03 | 21,438,272 | 654 | 34.7% | 17.6% | 43.1% |
| Scaf_04 | 20,712,704 | 613 | 35.2% | 16.9% | 44.7% |
| Scaf_05 | 20,149,410 | 580 | 35.3% | 16.0% | 43.6% |
| Scaf_06 | 19,848,268 | 632 | 35.4% | 16.8% | 43.4% |
| Scaf_07 | 19,643,986 | 510 | 34.7% | 16.3% | 42.5% |
| Scaf_08 | 18,176,930 | 552 | 35.0% | 17.3% | 41.4% |
| Scaf_09 | 17,506,527 | 606 | 34.9% | 17.5% | 41.1% |
| Scaf_10 | 17,186,144 | 427 | 35.0% | 19.2% | 42.8% |
| Scaf_11 | 16,482,901 | 535 | 35.2% | 18.1% | 44.2% |
| Scaf_12 | 15,131,411 | 462 | 35.4% | 17.6% | 45.5% |
| Scaf_13 | 13,599,833 | 355 | 34.8% | 15.2% | 48.2% |
| Scaf_14 | 12,890,779 | 421 | 35.2% | 17.6% | 44.1% |
| Scaf_15 | 12,471,685 | 414 | 35.4% | 16.2% | 40.1% |
| Scaf_16 | 12,249,352 | 388 | 35.9% | 13.6% | 43.8% |
| Scaf_17 | 11,871,507 | 384 | 35.7% | 17.5% | 44.9% |
| Scaf_18 | 9,236,171 | 347 | 34.7% | 18.9% | 42.1% |
| Scaf_19 | 9,108,583 | 268 | 35.9% | 14.7% | 46.0% |
| Scaf_20 | 7,025,367 | 202 | 36.2% | 20.9% | 49.2% |
| Scaf_X | 20,422,211 | 663 | 35.2% | 12.2% | 40.7% |
| unincorporated | 12,174,703 | 2,263 | 35.8% | 17.0% | 42.1% |
| mtDNA | 18,007 | 13 | 23.9% | 0% | * |

Closer examination of genetic differences among individuals revealed that four non-synonymous (amino-acid changing) substitutions occurred in protein coding genes, and all of these were unique to single individuals. Other differences were found either at synonymous sites within protein coding genes or in RNA encoding regions (Fig. 4A). Two substitutions occurred at the tRNA^Leu^ and tRNA^Lys^ regions that bracket the protein-coding *cytochrome oxidase II* gene. These substitutions co-occur with three synonymous substitutions within *cytochrome oxidase II*, which were used as RFLP markers of different populations in prior studies (Dahlhoff et al. 2019; Rank et al. 2020). Two more substitutions occurred in the large ribosomal subunit (16S) region, and beetles with southern haplotypes (4 BP and 2 BC) possessed a two base pair insertion in this region. No substitutions occurred in regions of the mitochondrial genome where overlapping genes are found.

Analysis of putative secondary structure of tRNA and rRNA molecules produced by these substitutions revealed that nucleotide differences between northern and southern haplotypes are located in regions that could affect functional properties of the molecules. Within the 65 bp tRNA^Leu^ region, northern beetles possess a ‘C’ nucleotide at position 34, while southern beetles possess an ‘A’ nucleotide at the same position. This substitution is located in the loop portion of the anticodon, two base pairs away from the ‘TAA’ anticodon (Fig. S2). Within the 69 base pair tRNA^Lys^ region, the substitution from ‘T’ in the northern populations to ‘C’ in the southern ones occurs at position 37, which occurs in the final base in the stem portion of the anticodon and results in non-canonical base pairing, which would affect stability of a critical region within the molecule (Fig. S3). Finally, within the 16S ribosomal RNA region, two substitutions and one insertion distinguish northern and southern beetles, and these sequence differences result in differences in predicted secondary structure and free energy values (Fig. S3).

## DISCUSSION

Using a combination of 10x genomics sequencing, Hi-C scaffolding, transcriptomics, and population-level whole genome resequencing, we were able to assemble highly quality nuclear and mitochondrial genomes for the willow leaf beetle, *C. aeneicollis*. The total length of our nuclear assembly (356 Mb) is in the range of other chromosome-length beetle genome assemblies and provides an important resource for this underrepresented order of insects. Our annotation of 12,586 genes is also consistent with other high quality beetle genome assemblies (Fallon et al. 2018; Keeling et al. 2022; Van Dam et al. 2021), and by establishing orthology with the model species, *Tribolium castaneum* (Herndon et al. 2020) we have putative gene functions for over 60% of the genes in our annotation. The putative number of linkage groups proposed here (21) is roughly consistent with cytological studies of other *Chrysomela* (Petitpierre 2011). Within Chrysomelini, the ancestral haploid chromosome number appears to be 17. Our genome assembly suggests a genome size that is also consistent with earlier cytological investigations (Petitpierre 2011).

Mitochondrial differences along the latitudinal gradient show substantial differences between northern and southern populations and are consistent with nuclear genetic data suggesting that Bishop Creek populations reflect introgression between north and south (Rank et al. 2020). The number of substitutions separating Rock Creek and Big Pine mitotypes suggest that they have been distinct for a considerable length of time. Levels of differentiation are greater at mitochondrial than at nuclear loci in *C. aeneicollis* (Dellicour et al. 2014), which is consistent with predictions that mitochondria should evolve faster than the nuclear genome due to higher mutation rates (Burton and Barreto 2012), reduced effective population sizes (Neiman and Taylor 2009), and potential functional differences among mitotypes (Hill 2015) (although see Cooper et al. 2015; Edwards et al. 2021).

In *C. aeneicollis*, mitochondrial variation appears to be partly a result of natural selection and adaptation to local environmental conditions (Dahlhoff et al. 2019; Rank et al. 2020). Specifically, beetles are found at higher elevations in Big Pine Creek in the south than in Rock Creek, and larvae with the Big Pine Creek mitotype developed twice as fast as those with the northern mitochondrial haplotype when reared at 3800 m ASL at the Barcroft Station of the White Mountain Research Center (Dahlhoff et al. 2019). Conversely, when larvae were reared at 1240 m at the Owens Valley Station, running speed of beetles with northern mitotypes recovered from a non-lethal heat exposure more effectively than those with the southern mitotype (Rank et al. 2020). In addition, beetles with matching mitochondrial and nuclear genetic backgrounds (southern/southern or northern/northern) show higher reproductive success and faster larval development in the field (Rank et al. 2020).

Our results show that most differences between northern and southern mitotypes occur at synonymous sites within protein coding genes, and all non-synonymous substitutions were unique to a single beetle. In contrast, differences between northern and southern mitotypes at two tRNA loci and the large ribosomal subunit are predicted to generate functional differences in secondary structure. These differences could result in incompatibilities between nuclear-encoded genes that function inside mitochondria to perform protein synthesis and their mitochondrial RNA counterparts (Hoekstra et al. 2013). Our results provide a potential mechanism for the mitonuclear epistasis described in prior studies (Rank et al. 2020) as three critical metabolic processes require integration between mitochondrial and nuclear gene products (Hill 2015; 2016; Hill et al. 2019; Levin et al. 2014). These processes include interactions among proteins involved in the electron transport chain, mitochondrial replication inside cells, and protein synthesis inside the mitochondrion (Levin et al. 2014). This third mechanism has been proposed to operate in lab-generated lines of Drosophila that were created to generate mitochondrial mismatch (Hoekstra et al. 2013).

In summary, the genomic information presented in this paper adds important resources that expand our understanding of beetle genomics and will help those studying the genetics and genomics of host plant use in phytophagous insects. Coupled with our mtDNA assembly, these resources will be of great use to those investigating the evolutionary physiology of ectotherm metabolism in natural environments characterized by multiple axes of abiotic stress.

## ACKNOWLEDGEMENTS

We thank A. Lumley, who oversaw larval rearing experiments and completed sample preparation and DNA extraction, and B. Sargeant, M. Smith, V.C. Dahlhoff, I. Erundu, E. Lopez, D. Martinez, R. Regello, and R. Bravo for assistance in the field. Members of the Lanzaro laboratory at UC-Davis, especially Y. Lee, helped with DNA extraction. We thank P. Mardulyn, C. Wheat and E. Hornett for help generating the initial draft mitochondrial genome and J. Deyarmin for work on mitochondrial gene products. This project was supported by grants from the National Science Foundation IOS 1558159 to C. M. Williams and J.H. Stillman, IOS 1457335 to N. E. Rank, and IOS 14573 to E.P. Dahlhoff, a California Conservation Genomics grant to C.M. Williams, D. Bachtrog, N.E. Rank, and E.P. Dahlhoff, and computing resources through XSEDE Allocation IBN180012 to C.M. Williams and J.H. Stillman.

## DATA ACCESSIBILITY STATEMENT

All data used in the genome assembly and annotation are described and available under bioproject: PRJNA693623. The genome assembly accession at NCBI is GCA_027562985.1 and the mtDNA accession is OPT787486. All associated files are also available here: https://figshare.com/projects/A_chromosome_scale_genome_assembly_and_evaluation_of_mtDNA_variation_in_the_willow_leaf_beetle_Chrysomela_aeneicollis/152286

## SUPPLEMENTAL FIGURE CAPTIONS

**Figure S1**. A) Karyoplot of 21 linkage groups and the proportion of masked repetitive sequence along the genome (blue, 50 kb windows) with gene density (high to low) shown as a white:green:black heatmap in 500 kb genomic intervals. B) Repetitive sequence distributions across the genome assembly plotted as the percentage of bases masked in 50 kb windows per repetitive sequence type. Regions found to be enriched for repeats (defined as at least two consecutive windows ≥ 2× the genome-wide average) highlighted with red boxes.

**Figure S2**. A) Region bracketing the cytochrome oxidase gene with tRNA molecules that differ between Big Pine Creek and Rock Creek. B) Predicted secondary structure of the tRNA^Leu^ alleles that differ between Big Pine Creek and Rock Creek. C) Predicted secondary structure of the tRNA^Lys^ alleles that differ between Big Pine Creek and Rock Creek.

**Figure S3**. A) Predicted secondary structure of the large ribosomal subunit gene for individuals possessing the northern mitochondrial genotype found in Rock Creek. B) Predicted secondary structure of the large ribosomal subunit gene for individuals possessing the mitochondrial genotype found in Big Pine Creek.

